# Calibrated Uncertainty Quantification for Patient-Level AML Drug Sensitivity Prediction Using Split Conformal Prediction

**DOI:** 10.64898/2026.06.07.730728

**Authors:** Amir J. Shokrzadeh, Pegah Shokrzadeh

## Abstract

Accurate prediction of ex vivo drug sensitivity in acute myeloid leukemia (AML) patients from transcriptomic data is a critical challenge for precision oncology. Existing computational approaches have explored uncertainty quantification in cancer drug response prediction primarily using cell line data, while patient-level AML models typically rely on heuristic confidence measures rather than statistically calibrated uncertainty estimates. Here, we present a framework applying split conformal prediction to patient-level AML drug response modeling using the BeatAML 2.0 cohort. We trained Elastic Net and XGBoost regressors on bulk RNA-seq gene expression profiles from 318 AML patients, analyzing 34,764 patient-drug observations across 122 compounds. Baseline models achieved median Pearson R values of 0.291 (Elastic Net) and 0.281 (XGBoost) across 122 drugs. Wrapping these models with split conformal prediction yielded well-calibrated prediction intervals across three confidence levels: empirical coverages of 81.4%, 90.7%, and 95.5% against nominal targets of 80%, 90%, and 95%, respectively. Analysis of prediction interval widths revealed substantial drug-class-specific uncertainty patterns, with HDAC and BCL-2 inhibitors exhibiting markedly higher uncertainty than MDM2 inhibitors, suggesting a potential association between transcriptomic predictability and drug mechanism of action, although several drug classes were represented by only a small number of compounds. Predictive uncertainty was not significantly associated with ELN2017 molecular risk classification (Kruskal-Wallis p=0.395) or NPM1 mutation status (p=0.788). These results demonstrate that statistically valid uncertainty quantification can be achieved for patient-level AML drug response prediction despite substantial biological heterogeneity. to the best of our knowledge, no published study has applied split conformal prediction to patient-level ex vivo drug sensitivity prediction in the BeatAML cohort, providing a principled alternative to heuristic confidence scoring approaches.

## Introduction

Acute myeloid leukemia (AML) is an aggressive hematologic malignancy characterized by clonal expansion of myeloid progenitor cells and marked inter-patient heterogeneity in treatment response^1^. Despite advances in targeted therapy, treatment selection remains largely empirical, with many patients failing to respond to initial regimens^2^. Ex vivo drug sensitivity profiling, measuring the response of patient-derived leukemic cells to a panel of compounds, offers a direct functional readout of individual drug susceptibility, providing a promising basis for treatment personalization^3,2^.

Computational prediction of ex vivo drug sensitivity from molecular profiles has emerged as a complementary approach to functional screening, enabling predictions for patients where fresh sample availability is limited. The BeatAML project represents the largest publicly available patient-level resource linking bulk RNA sequencing, whole-exome sequencing, and ex vivo drug sensitivity data across hundreds of AML patients^3,2^. Machine learning methods applied to this dataset include Elastic Net regression integrating RNA sequencing, mutation, and clinical features^5^, network-driven frameworks integrating pathway information^6^, and MDREAM, a multi-omics integration approach using Support Vector Machines that achieved Spearman correlations up to 0.68 in cross-cohort validation^4^. Despite this progress, predictive performance remains moderate across the full drug panel, reflecting the substantial transcriptomic heterogeneity of AML patient samples.

A critical limitation of existing patient-level drug response models is their reliance on point predictions without statistically calibrated uncertainty estimates. MDREAM incorporates a heuristic confidence score to indicate prediction reliability^4^; while practically useful, heuristic confidence scores lack formal statistical coverage guarantees — they do not ensure that a stated confidence level corresponds to a verified empirical probability of containing the true response value. This distinction is clinically meaningful: a model reporting high confidence without calibration verification may systematically mislead treatment decisions in a high-stakes oncology setting.

Conformal prediction provides a distribution-free framework for constructing prediction intervals with guaranteed marginal coverage under the assumption of exchangeable data^7,8^. As with all conformal methods, coverage guarantees rely on the assumption that calibration and test observations are exchangeable. Split conformal prediction achieves this by partitioning data into training, calibration, and test sets, using held-out calibration residuals to set interval widths that provably cover the true outcome at a user-specified rate^9^. Recent work has applied conformal prediction to anti-cancer drug sensitivity prediction using cell line data from the Genomics of Drug Sensitivity in Cancer (GDSC) database^10,11^, demonstrating valid calibration and improved reliability over point prediction approaches. However, to the best of our knowledge, no published study has applied split conformal prediction to patient-level ex vivo drug sensitivity prediction using the BeatAML cohort.

Here, we address this gap by presenting a framework for calibrated uncertainty quantification in AML drug response prediction. We apply split conformal prediction as a wrapper around Elastic Net and XGBoost regressors trained on bulk RNA-seq data from 318 AML patients across 122 drugs in the BeatAML 2.0 cohort. We evaluate empirical coverage across multiple confidence levels and characterize uncertainty patterns across drugs and drug classes. We further explore whether drug-level uncertainty patterns show associations with established AML molecular subgroup classifications. Our results demonstrate that statistically valid prediction intervals are achievable for patient-level AML drug response modeling, and suggest that predictive uncertainty may be more strongly associated with drug mechanism of action than with standard AML molecular subgroup classifications.

## Methods

### Dataset

We utilized data from the BeatAML 2.0 study (Bottomly et al., Cancer Cell 2022), accessed via cBioPortal and the BeatAML GitHub repository. Gene expression data consisted of mRNA z-scores computed relative to all samples (log2 RPKM), available for 671 specimens across 562 patients. Ex vivo drug sensitivity data were obtained from Supplementary Table S10 of the original publication, containing AUC values for 122 compounds across 528 patients identified by internal laboratory identifiers. Clinical and mutation annotations were obtained from the cBioPortal data package, including ELN2017 risk classification, FLT3-ITD status, and NPM1 mutation status. Patient identifier mapping between cBioPortal sample IDs and original laboratory identifiers was performed using the BeatAML sample mapping file (beataml_waves1to4_sample_mapping.xlsx) from the BeatAML GitHub repository.

### Cohort Construction

Gene expression samples were transposed to a patient × gene matrix. Where multiple specimens were available for a single patient, expression values were averaged to yield one observation per patient. Nine genes with missing values across all patients were removed prior to analysis. Variance filtering was applied to retain genes with variance greater than 1.0 across the cohort, a threshold selected empirically based on inspection of the variance distribution to reduce dimensionality while retaining highly variable genes; no formal sensitivity analysis of the variance threshold was performed. This reduced the feature set from 22,834 to 14,121 genes. Drug sensitivity records were linked to gene expression profiles via the sample mapping file. Only patients with both transcriptomic profiles and drug sensitivity measurements were retained, yielding a modellable cohort of 318 patients with matched data across 122 drugs, comprising 34,764 patient-drug observations. For clinical annotation, one sample per patient was retained by prioritizing specimens marked as used in the manuscript. ELN2017 categories were harmonized to four groups (Favorable, Intermediate, Adverse, NonInitial); ambiguous or rare labels (n=16 patients) were excluded from subgroup analyses only, not from model training or evaluation.

### Baseline Prediction Models

For each of the 122 drugs independently, two regression models were trained to predict AUC from gene expression features: Elastic Net (alpha=0.1, l1_ratio=0.5, max_iter=5000; scikit-learn) and XGBoost (n_estimators=100, max_depth=3, learning_rate=0.1, subsample=0.8, colsample_bytree=0.3). Hyperparameters were fixed a priori to focus the study on the uncertainty quantification framework rather than predictive model optimization. For each drug, available patient-drug pairs were split into 80% training and 20% test sets (random_state=42). No drugs met the exclusion threshold of fewer than 30 patients. Model performance was evaluated on the held-out test set using Pearson correlation coefficient (R) and root mean squared error (RMSE); median values across all 122 drugs are reported.

### Conformal Prediction

Split conformal prediction was applied to the Elastic Net models using the SplitConformalRegressor class from MAPIE (v1.0.1) with prefit=True. For each drug, the 80% training portion was further partitioned into 60% model training and 20% calibration, yielding approximate splits of 60% train, 20% calibration, and 20% test relative to the full per-drug sample. The fitted Elastic Net model was conformalized using the held-out calibration set via the conformalize() method. Prediction intervals were generated at three nominal confidence levels: 80%, 90%, and 95%. Empirical coverage was first computed independently for each drug as the proportion of test patients whose true AUC fell within the predicted interval, then averaged across drugs (macro-average), with each drug contributing equally regardless of test set size. Mean interval width was computed as the mean difference between upper and lower interval bounds across test patients per drug. Drugs with insufficient calibration samples for a given confidence level (minimum required: ⌈1/(1−α)⌉ + 1 samples) were excluded from coverage computation at that level. Coverage was computed across all 122 drugs at the 80% and 90% levels, and across 114 drugs at the 95% level, as 8 drugs had fewer than 20 calibration samples. All analyses were performed on CPU hardware.

### Uncertainty Analysis

Prediction interval widths at the 90% confidence level were used as the primary measure of predictive uncertainty. Because split conformal prediction produces a single calibration quantile per drug, interval widths are uniform within each drug and reflect drug-level rather than patient-level uncertainty. Mean interval width was compared across drug classes defined by mechanism of action: BCL-2 inhibitors, HDAC inhibitors, FLT3 inhibitors, MEK inhibitors, mTOR inhibitors, Hedgehog pathway inhibitors, and MDM2 inhibitors. For molecular subgroup analyses, prediction interval widths from all patient-drug test observations were grouped by patient molecular annotation. Observations from patients belonging to each ELN2017 risk group were compared using the Kruskal-Wallis test; FLT3-ITD and NPM1 mutation status groups were compared using the Mann-Whitney U test (two-sided). Each patient contributed multiple observations corresponding to the drugs in which they appeared in the test set. Effect sizes for pairwise comparisons were computed as rank-biserial correlation. All statistical analyses were performed using scipy.stats (Python).

### Reproducibility

All code is available at https://github.com/Amir-Shokrzadeh/aml-conformal-prediction(v1.0). Data are publicly available via cBioPortal (https://www.cbioportal.org) and the BeatAML GitHub repository (https://github.com/biodev/beataml2.0_data).

## Results

### Baseline Drug Response Prediction Performance

Elastic Net and XGBoost regressors were trained independently for each of the 122 drugs in the BeatAML 2.0 cohort. Across all drugs, Elastic Net achieved a median Pearson R of 0.291 (interquartile range: 0.171–0.391) and median RMSE of 46.7 AUC units. XGBoost achieved comparable performance with a median Pearson R of 0.281 (IQR: 0.184–0.400) and median RMSE of 43.4 AUC units. Performance varied substantially across drugs: 28 drugs achieved Pearson R > 0.4 with Elastic Net, while 5 drugs showed negative correlation, indicating that transcriptomic features were uninformative for those compounds. The highest-performing drugs included Sorafenib (XGBoost R=0.623), Trametinib (R=0.607), and 17-AAG (R=0.588). These results are consistent with prior reports of moderate predictive performance for patient-level AML drug response prediction using transcriptomic data alone (Figure 1).

**Fig 1.**
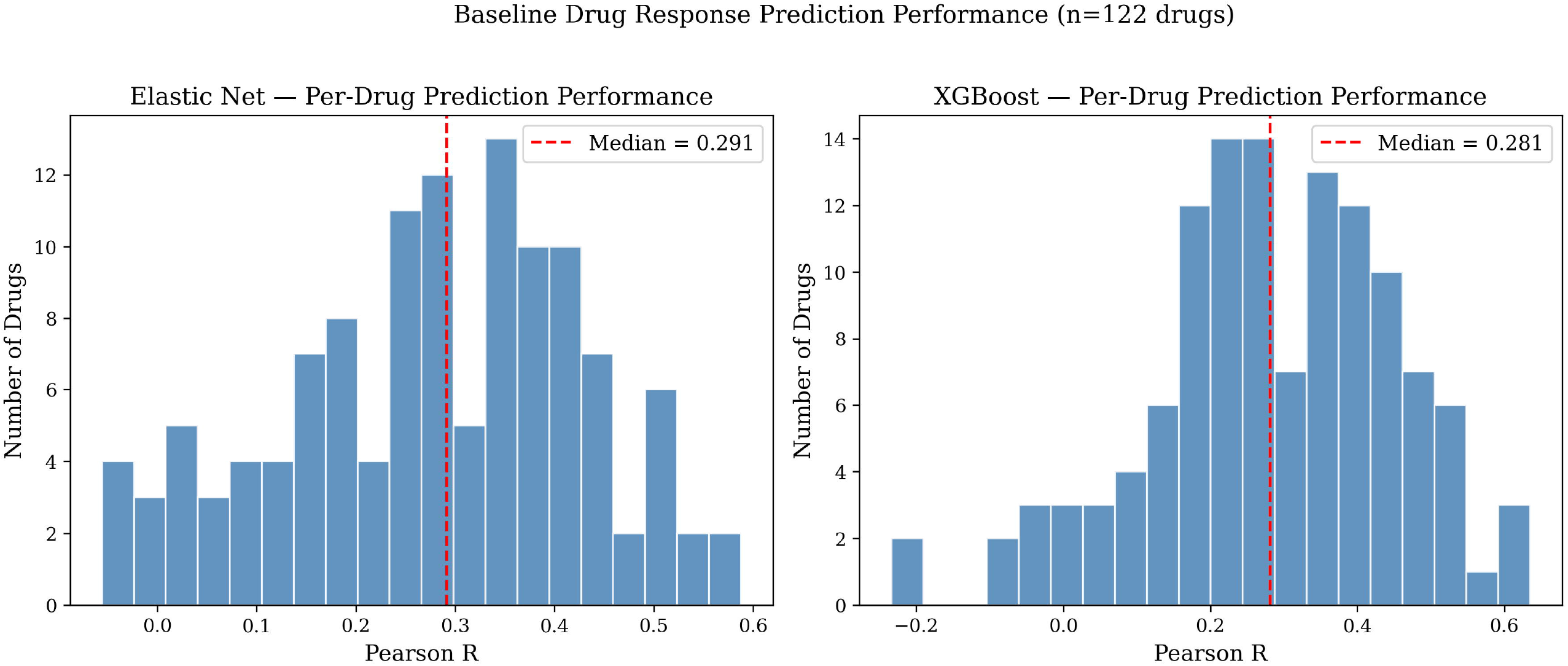
Baseline drug response prediction performance across 122 AML drugs. Distribution of Pearson correlation coefficients (R) between predicted and observed AUC values for Elastic Net (left) and XGBoost (right) regressors, evaluated on held-out test sets. Each histogram represents one drug model trained on bulk RNA-seq gene expression data from the BeatAML 2.0 cohort (318 patients). Dashed red lines indicate median Pearson R across all 122 drugs (Elastic Net: 0.291; XGBoost: 0.281).

### Conformal Prediction Calibration

Split conformal prediction was applied to the Elastic Net models for all 122 drugs at three nominal confidence levels. Empirical coverage closely matched nominal targets across all levels: 81.4% at the 80% level (n=122 drugs), 90.7% at the 90% level (n=122 drugs), and 95.5% at the 95% level (n=114 drugs; 8 drugs excluded due to insufficient calibration samples). These results indicate that the split conformal prediction framework produced well-calibrated prediction intervals in the BeatAML cohort. Calibration curves for individual drugs showed variability around the diagonal, particularly for drugs with small test sets, but mean empirical coverage across drugs closely tracked nominal levels at all three confidence targets (Figure 2). Mean prediction interval widths at the 90% confidence level ranged from 77.5 to 272.1 AUC units across drugs (overall mean: 162.4 AUC units), reflecting substantial variation in predictive uncertainty across compounds.

**Fig 2.**
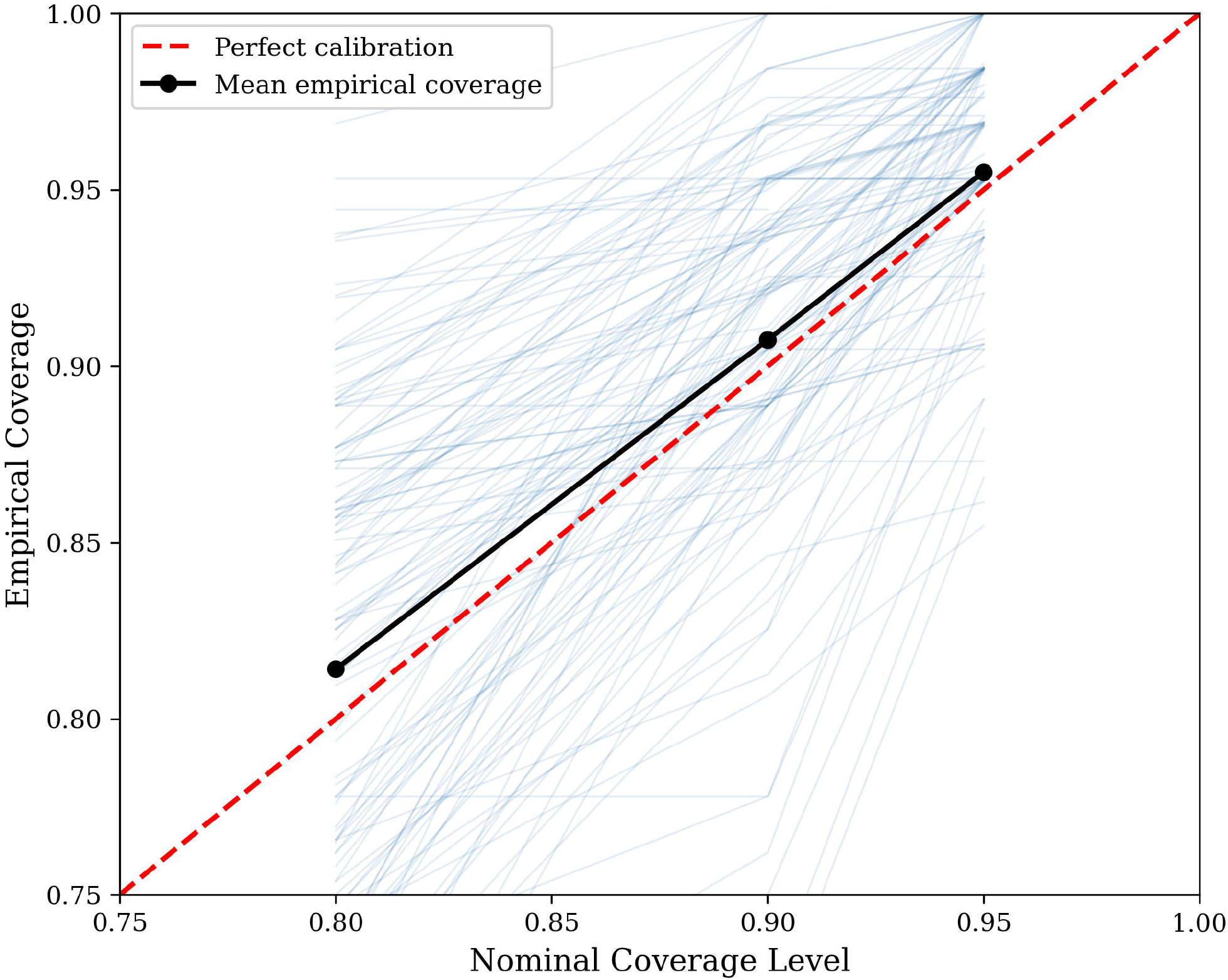
Conformal prediction calibration across 122 AML drugs. Empirical versus nominal coverage for split conformal prediction intervals at three confidence levels (80%, 90%, 95%). Each blue line represents one drug (n=122 at 80% and 90%; n=114 at 95%). The red dashed diagonal indicates perfect calibration. The black line with filled circles indicates mean empirical coverage across drugs at each nominal level (80%: 81.4%; 90%: 90.7%; 95%: 95.5%). Coverage was computed as a macro-average across drugs, with each drug contributing equally regardless of test set size.

### Drug-Class Uncertainty Patterns

Analysis of prediction interval widths at the 90% confidence level revealed substantial variation in predictive uncertainty across drug classes defined by mechanism of action (Figure 3). HDAC inhibitors exhibited the highest mean uncertainty (260.3 AUC units; n=1 drug: Panobinostat), followed by BCL-2 inhibitors (242.9 AUC units; n=2 drugs: Venetoclax, ABT-737), FLT3 inhibitors (193.7 AUC units), and MEK inhibitors (192.7 AUC units). mTOR inhibitors showed intermediate uncertainty (157.6 AUC units), while Hedgehog pathway inhibitors (89.0 AUC units) and MDM2 inhibitors (77.5 AUC units) exhibited the lowest uncertainty. At the individual drug level, Venetoclax had the highest mean interval width (272.1 AUC units) and Nutlin 3a the lowest (77.5 AUC units) across all 122 compounds (Figure 4). This approximately 3.4-fold range in interval width highlights substantial differences in predictive uncertainty across drug classes.

**Fig 3.**
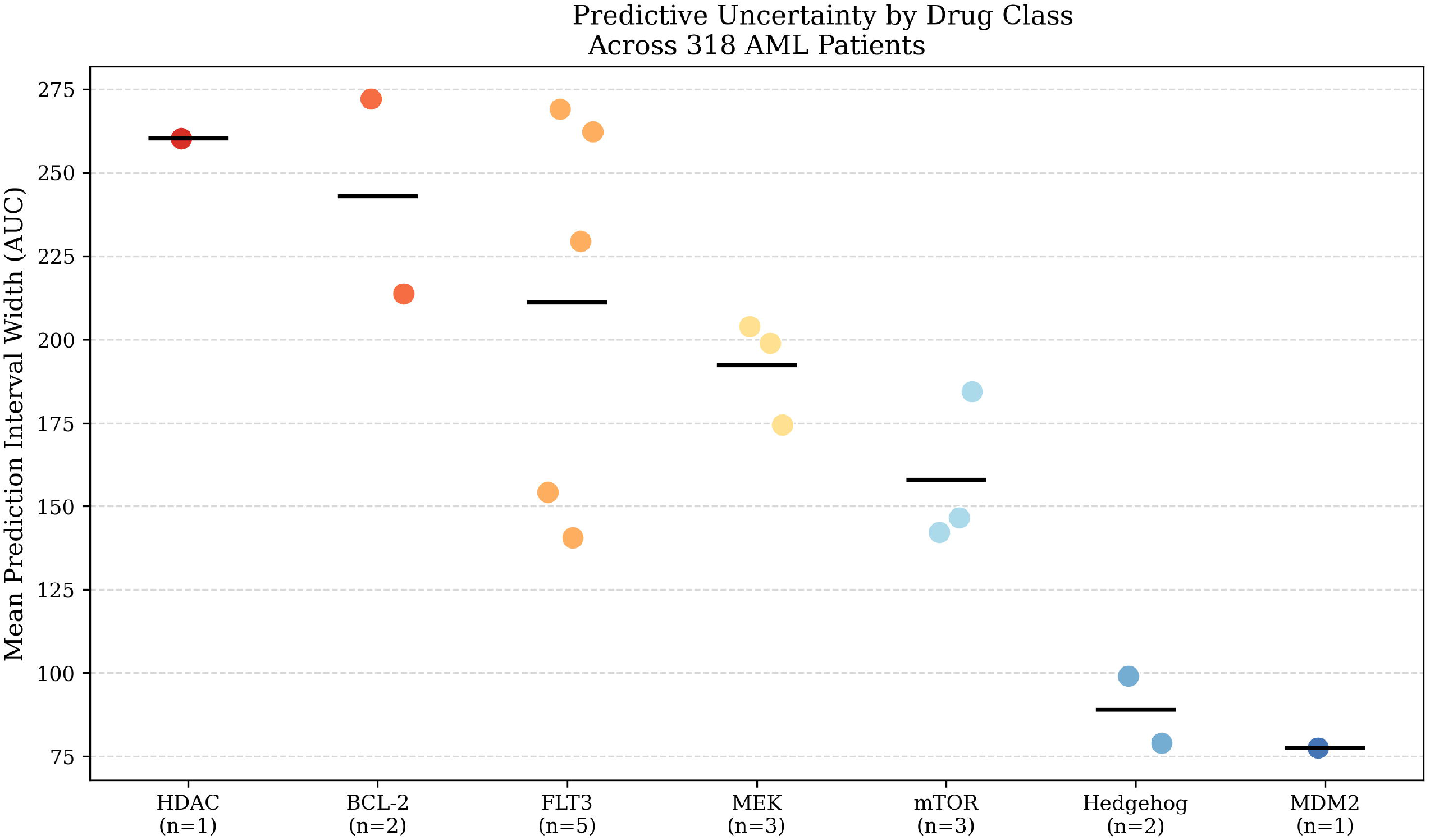
Predictive uncertainty by drug mechanism of action. Mean prediction interval width at the 90% confidence level for individual drugs grouped by mechanism of action class. Each dot represents one drug; horizontal black lines indicate the class mean. Drug classes are ordered by decreasing class mean. The number of drugs per class is shown in parentheses on the x-axis. Classes represented by a single drug, HDAC (Panobinostat) and MDM2 (Nutlin 3a), reflect individual drug characteristics rather than class-level patterns and should be interpreted accordingly.

**Fig 4.**
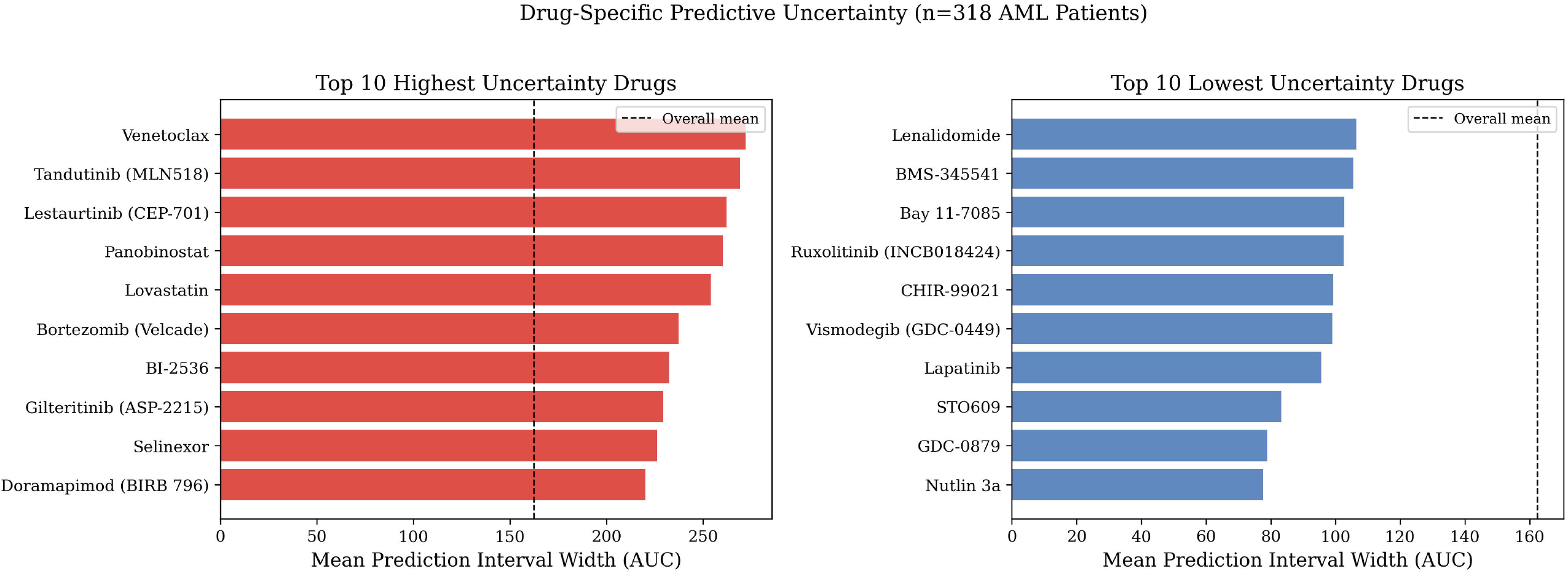
Drug-specific predictive uncertainty across 122 AML compounds. Mean prediction interval width at the 90% confidence level for the ten highest-uncertainty drugs (left, red) and ten lowest-uncertainty drugs (right, blue). The dashed vertical line indicates the overall mean interval width across all 122 drugs (162.4 AUC units). Drugs are ordered by mean interval width within each panel.

### Molecular Subgroup Uncertainty Analysis

Predictive uncertainty was examined in relation to established AML molecular subgroup classifications. No significant difference in prediction interval width was observed across ELN2017 risk groups (Favorable, Intermediate, Adverse, NonInitial; Kruskal-Wallis H=2.98, p=0.395), indicating that ELN2017 molecular risk classification does not predict transcriptomic uncertainty. FLT3-ITD mutation status showed a statistically significant but negligible association with interval width (Mann-Whitney U, p=0.014; rank-biserial r=0.038, mean difference 2.1 AUC units), with FLT3-ITD positive patients showing marginally wider intervals (159.9 ± 36.9 AUC units) compared to FLT3-ITD negative patients (157.8 ± 36.9 AUC units). Given the negligible effect size, this difference is not considered clinically meaningful. NPM1 mutation status showed no significant association with interval width (p=0.788). Taken together, these results indicate that predictive uncertainty at the drug-observation level is not meaningfully stratified by standard AML molecular subgroup classifications. Because split conformal prediction produces uniform interval widths within each drug by construction, all variation in interval width across the cohort is attributable to drug identity. The heatmap of interval widths across selected drugs and patients illustrates this drug-driven structure across the patient cohort (Figure 5).

**Fig 5.**
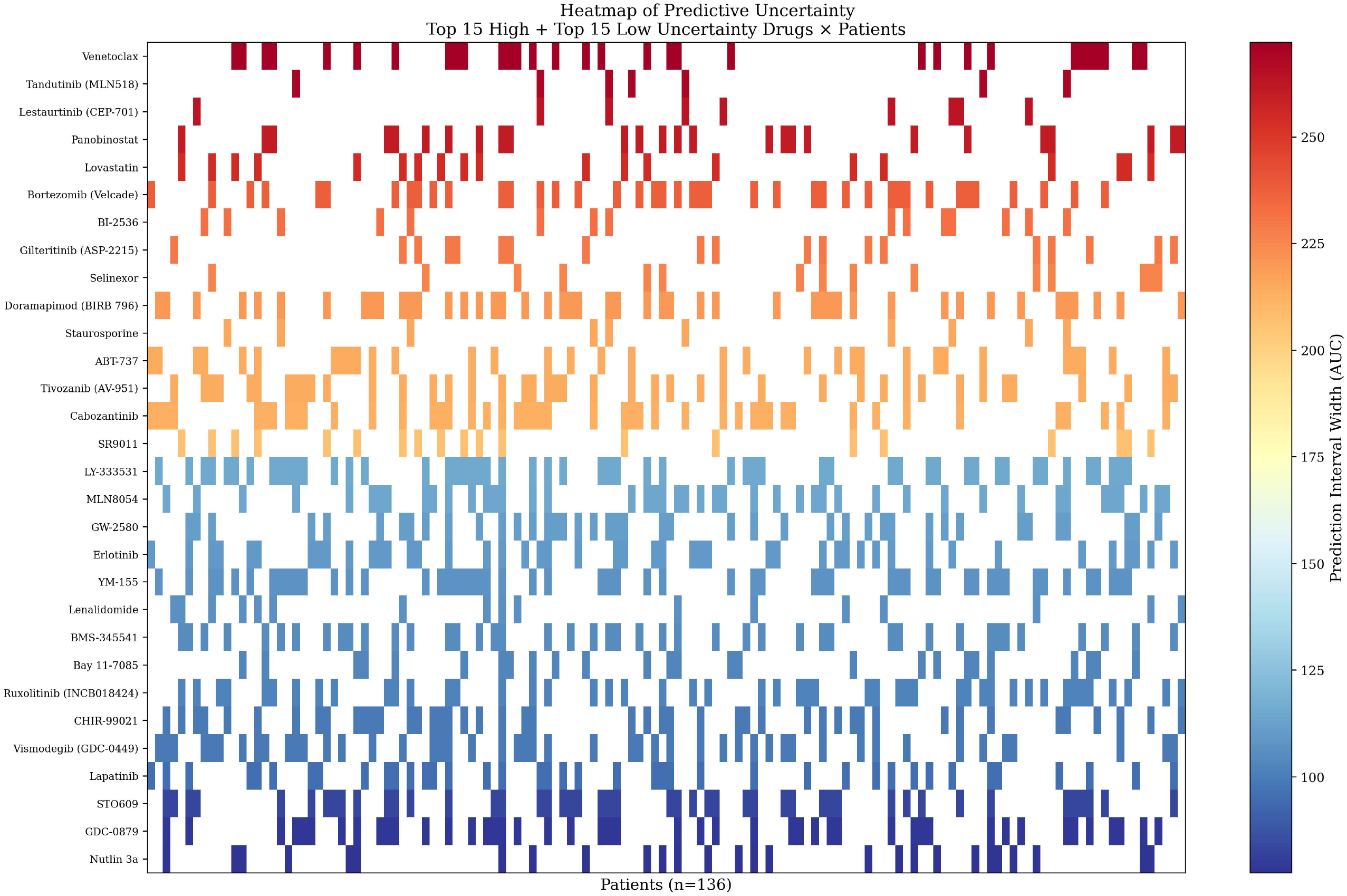
Heatmap of predictive uncertainty across drugs and patients. Prediction interval widths at the 90% confidence level for the 15 highest-uncertainty and 15 lowest-uncertainty drugs (rows) across 136 patients appearing in the test sets of at least 5 of the 30 selected drugs (columns). Color indicates interval width (AUC units); warmer colors indicate higher uncertainty. White cells indicate patients not included in the test set for that drug, reflecting the split conformal prediction design rather than missing data. Because split conformal prediction produces uniform interval widths within each drug by construction, all color variation across rows reflects drug identity rather than patient-specific uncertainty.

## Discussion

### Principal Findings

In this study, we applied split conformal prediction to patient-level ex vivo AML drug response prediction using the BeatAML 2.0 cohort, demonstrating that statistically calibrated uncertainty quantification is achievable across 122 drugs and 318 patients. Empirical coverage closely matched nominal targets at all three confidence levels tested, indicating that the framework produced well-calibrated prediction intervals in the BeatAML cohort. To the best of our knowledge, this represents the first application of split conformal prediction to patient-level ex vivo drug sensitivity prediction in the BeatAML cohort, addressing a gap left by prior approaches that rely on heuristic confidence scoring without formal coverage guarantees.

### Baseline Prediction Performance in Context

Baseline models achieved median Pearson R of 0.291 (Elastic Net) and 0.281 (XGBoost) across 122 drugs, consistent with prior reports of moderate predictive performance for patient-level AML drug response prediction using transcriptomic data alone^4,2^. This performance level reflects the substantial biological heterogeneity inherent to AML patient cohorts — inter-patient variation in clonal composition, co-occurring mutations, and microenvironmental factors contributes noise that bulk RNA-seq alone cannot fully capture. Our cohort of 318 patients with matched transcriptomic and drug sensitivity data is comparable in scale to prior AML drug response modeling efforts on BeatAML; MDREAM, for instance, used 278 patients for training and 461 total BeatAML patients across training and validation^4^, while our analysis required simultaneous availability of both gene expression profiles and drug sensitivity measurements, which reduced the modellable cohort from the full 562-patient dataset. The moderate predictive performance observed here does not diminish the value of the conformal prediction framework; on the contrary, it underscores the clinical necessity of calibrated uncertainty estimates, as point predictions from models with R≈0.29 carry inherent uncertainty that practitioners need to quantify rather than ignore.

### Calibration and the Advantage Over Heuristic Confidence Scoring

The central methodological contribution of this work is the demonstration that split conformal prediction produces well-calibrated prediction intervals for patient-level AML drug response data. Empirical coverage of 81.4%, 90.7%, and 95.5% against nominal targets of 80%, 90%, and 95% respectively indicates that the framework achieved its theoretical guarantee in practice on this dataset. This is a meaningful advance over heuristic confidence scoring approaches such as that employed by MDREAM^4^, which provide a useful but statistically informal indication of prediction reliability. Heuristic scores are not guaranteed to correspond to verifiable coverage probabilities — a model reporting 90% confidence using a heuristic measure may empirically achieve 60% or 99% coverage, with no principled way to verify which. Split conformal prediction eliminates this ambiguity by construction, providing practitioners with intervals whose coverage properties are formally guaranteed under the exchangeability assumption. This guarantee holds regardless of the underlying model, making the framework applicable to any future improvement in AML drug response prediction models.

### Drug-Class Uncertainty Patterns and Biological Interpretation

A notable finding of this study is the substantial variation in predictive uncertainty across drug classes, with an approximately 3.4-fold difference in mean interval width between the highest-uncertainty class (HDAC inhibitors, 260.3 AUC units) and lowest-uncertainty class (MDM2 inhibitors, 77.5 AUC units). We offer the following biological interpretations as hypotheses rather than established findings, as our data do not permit causal conclusions.

One possible explanation for the observed uncertainty hierarchy is that drug classes differ in how closely their response determinants are captured by bulk RNA-seq profiles. For instance, BCL-2 inhibitor response is known to depend substantially on protein-level expression of BCL-2 family members and mitochondrial priming state^12^, factors not directly measured by mRNA abundance. Similarly, HDAC inhibitor response involves epigenetic reprogramming whose transcriptomic consequences are complex and context-dependent. In contrast, MDM2 and Hedgehog pathway inhibitors may act through mechanisms with relatively consistent transcriptomic correlates, potentially contributing to their lower uncertainty. These interpretations are speculative; our data do not permit causal conclusions, and experimental validation would be required to confirm whether these biological factors drive the observed uncertainty differences.

It should be noted that several drug classes in this analysis were represented by only one or two compounds, notably MDM2 inhibitors (n=1 drug: Nutlin 3a) and HDAC inhibitors (n=1 drug: Panobinostat); BCL-2 inhibitors were represented by two compounds (Venetoclax and ABT-737, mean width 242.9 AUC units). Conclusions about sparsely represented classes should therefore be interpreted with caution, as uncertainty estimates reflect individual drug characteristics rather than well-sampled class-level patterns. Classes with broader representation, such as FLT3 inhibitors (n=5) and MEK inhibitors (n=3), provide more robust class-level estimates.

### Absence of Molecular Subgroup Associations

The finding that predictive uncertainty was not significantly associated with ELN2017 risk classification (p=0.395) or NPM1 mutation status (p=0.788) warrants careful interpretation. ELN2017 classification is a prognostic tool based on cytogenetic and molecular features, designed to predict overall survival rather than ex vivo drug sensitivity. It is therefore not necessarily expected to predict transcriptomic model uncertainty. The absence of an association suggests that the difficulty of predicting drug response from transcriptomics is not simply a function of molecular risk category, but rather depends on drug-specific biological factors as discussed above. FLT3-ITD status showed a statistically significant but negligible association with interval width (p=0.014, rank-biserial r=0.038, mean difference 2.1 AUC units), which we attribute to the large number of observations (n=6,899) providing sufficient statistical power to detect clinically irrelevant differences. We report this finding for completeness but do not consider it biologically meaningful.

## Limitations

Several limitations of this study should be acknowledged. First, standard split conformal prediction produces constant prediction interval widths within each drug, as interval size is determined by a single nonconformity quantile derived from the calibration set. Consequently, the framework as implemented cannot distinguish uncertainty between individual patients receiving the same drug; all variation in interval width is attributable to drug identity rather than patient characteristics. Future work may explore normalized, locally adaptive, or Mondrian conformal prediction approaches capable of producing patient-specific interval widths, enabling a more granular characterization of predictive reliability.

Second, baseline model hyperparameters were fixed a priori and not optimized across the full drug panel. While this is appropriate given the study’s focus on uncertainty quantification rather than predictive model optimization, tuned models would likely yield narrower prediction intervals and improved coverage efficiency.

Third, eight of 122 drugs had fewer than 20 calibration samples, resulting in their exclusion from the 95% confidence level analysis. Results for these drugs should be interpreted with caution.

Fourth, the analysis relied exclusively on bulk RNA-seq data. Factors known to influence AML drug response, including protein expression levels, clonal heterogeneity, bone marrow microenvironment interactions, and epigenetic state, are not captured by transcriptomic profiles alone. Integration of multi-omics data — proteomics, ATAC-seq, or single-cell RNA-seq — may improve both predictive performance and uncertainty calibration in future work.

Fifth, mutation annotations for TP53, RUNX1, and ASXL1 were available for fewer than 30 patients in our modellable cohort, precluding subgroup analysis for these clinically important mutations.

Sixth, molecular subgroup analyses were performed on patient-drug observations rather than independent patients, as interval width is uniform within each drug by construction. Consequently, each patient contributes multiple non-independent observations. P-values from these analyses should therefore be interpreted as exploratory rather than confirmatory.

## Future Directions

The conformal prediction framework presented here is model-agnostic and can be applied to any regression model producing AUC predictions from molecular features. Natural extensions include application to deep learning architectures such as transformers or graph neural networks trained on multi-omics data, which may improve point prediction performance and consequently yield narrower, more clinically useful prediction intervals. The integration of Mondrian conformal prediction, which conditions interval construction on patient subgroup membership, could enable uncertainty estimates that are calibrated within molecular subtypes rather than marginally across the full cohort. Finally, prospective validation of this framework in an independent AML cohort would strengthen confidence in the generalizability of both the calibration results and the drug-class uncertainty patterns observed here.

## Conclusion

We presented a framework for calibrated uncertainty quantification in patient-level AML drug response prediction using split conformal prediction. Applied to 318 AML patients and 122 drugs from the BeatAML 2.0 cohort, the framework produced well-calibrated prediction intervals across three confidence levels, with empirical coverages of 81.4%, 90.7%, and 95.5% against nominal targets of 80%, 90%, and 95% respectively. These results indicate that statistically valid uncertainty quantification is achievable for patient-level ex vivo drug sensitivity prediction using transcriptomic data, without requiring distributional assumptions about the underlying data.

Analysis of prediction interval widths revealed substantial variation in predictive uncertainty across drug classes, with HDAC and BCL-2 inhibitors exhibiting the highest uncertainty and MDM2 and Hedgehog pathway inhibitors the lowest. This drug-class uncertainty hierarchy suggests that transcriptomic predictability may vary systematically by mechanism of action, potentially reflecting differences in how closely bulk mRNA expression captures the biological determinants of drug response for each class. In contrast, predictive uncertainty was not meaningfully associated with standard AML molecular subgroup classifications, including ELN2017 risk category and NPM1 mutation status.

The conformal prediction framework presented here is model-agnostic and readily extensible to more complex predictive architectures or multi-omics feature sets. By providing prediction intervals with formal coverage guarantees rather than heuristic confidence scores, this approach offers a principled foundation for assessing the reliability of computational drug response predictions in AML, a step toward more trustworthy computational decision-support tools in precision oncology.

